# A ‘comb’ algorithm for accurate detection of calcium transients and action potentials in regularly activating cardiac preparations

**DOI:** 10.1101/757294

**Authors:** Jakub Tomek

**Affiliations:** Department of Anatomy, Physiology, and Genetics, University of Oxford

## Abstract

Cardiac imaging and electrophysiological measurements yield vast amounts of data that typically need to be processed automatically. However, the detection and segmentation of calcium transients or action potentials is complicated by signal noise or signal drift, which may cause both false positive and negative segmentation. This article presents a simple but accurate ‘comb’ algorithm for detection of calcium transients and action potentials in such data where the pattern of activation is regular and its frequency is known. This corresponds either to cases where the cardiac preparation is paced externally, or where the preparation is beating in a stable rhythm. The prior knowledge of the heart rate is leveraged to overcome a broad range of artefacts and complications which arise in experimental data, such as different types of noise, signal drift, or alternans. The algorithm is simple to implement and has only a single free parameter, which is furthermore simple to set. A Matlab/Octave implementation of the comb algorithm is provided.

## 1 Algorithm description

The basic version of the presented algorithm is designed for detection of calcium transients in cardiac imaging data. It relies on finding the minima between adjacent calcium transients, reporting these as calcium transient boundaries. We first provide an informal graphical intuition of its principle when applied to calcium imaging. Second, the pseudocode is given. Third, we describe how the algorithm can be adapted for detection of action potentials in electrophysiological recordings or data from membrane potential imaging.

The code implementation of the algorithm (for Matlab and Octave) can be downloaded from the following GitHub address: https://github.com/jtmff/comb. This also includes a script which generates the figures used in this article.

### 1.1 Comb algorithm intuition for calcium imaging

The presented algorithm relies on prior knowledge of the *basic cycle length* (bcl) of the recording, i.e., how many ms apart are the starting times of calcium transients. The central visual intuition is that a ‘comb’ is positioned over the calcium fluorescence signal, with its teeth bcl ms apart^1^ (**Figure 1**). The algorithm works in two stages: 1) *comb positioning* and 2) *tooth refinement*.

**Figure 1:**
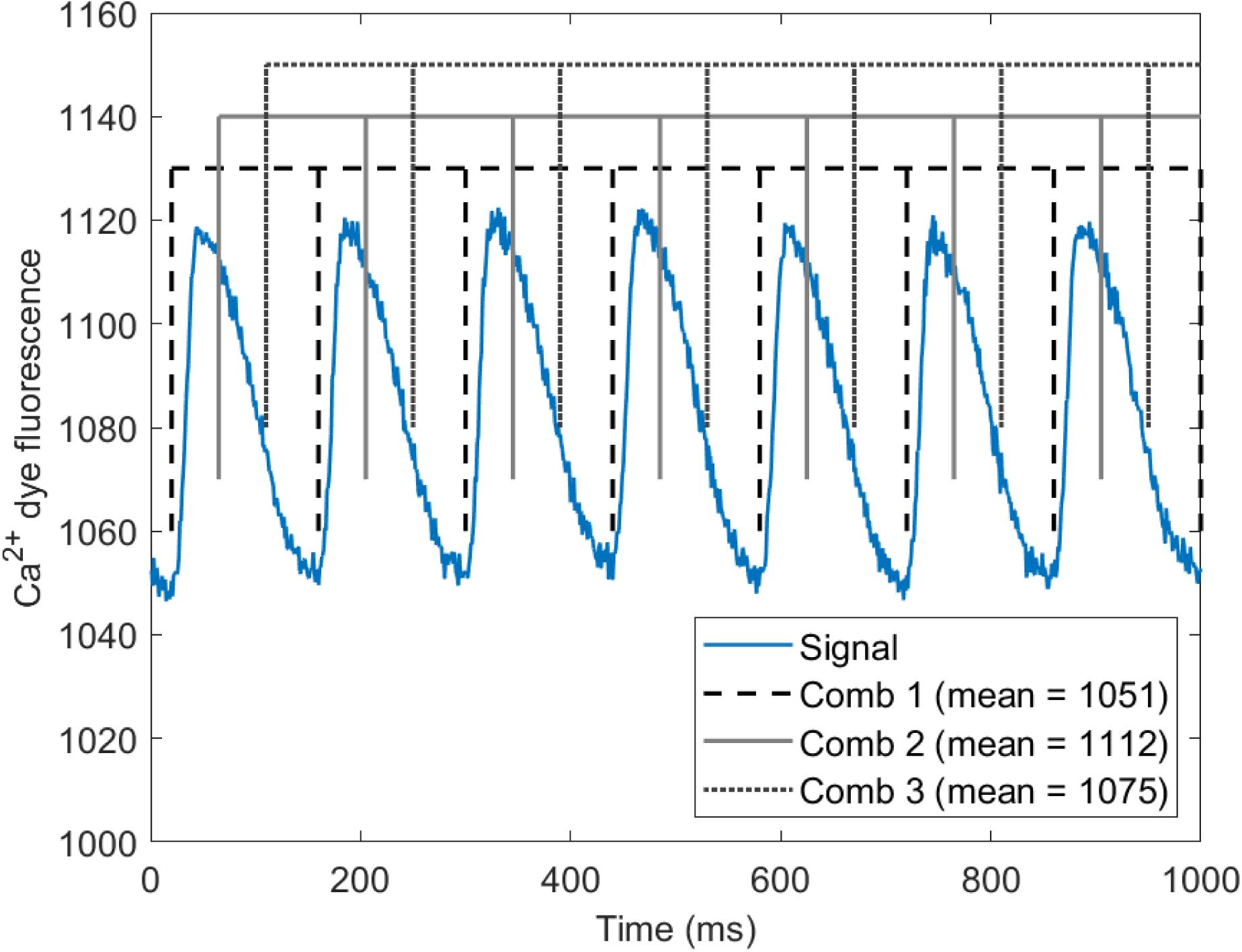
Comb algorithm illustration. An example of calcium imaging recording with basic cycle length of 140 ms, with three distinct positions of a ‘comb’. Means of values at x-axis locations corresponding to each comb’s teeth are given in the legend.

In *comb positioning*, the comb is slid over the signal, with the first tooth starting at 1 ms, 2 ms, …, bcl ms^2^. In each position, the mean of signal under the comb teeth is computed, and the minimum over these means is taken as the optimal comb position. Example of three comb positions and corresponding means under the teeth are given in **Figure 1**. The strength of this approach is that it utilizes information from the whole recording at once and the comb position with minimum average value under its teeth naturally corresponds to locations very close to signal minima.

In the second step, *tooth refinement*, each near-minimum identified as the location of one of the comb’s teeth is taken as a starting point of a local search. The search finds the minimum within *refinementWidth* ms around the comb tooth, where *refinementWidth* is a user-selected parameter. This step is designed for cases such as cardiac alternans or slightly irregular heartbeat, where the minima are not spaced fully equidistantly. The optimal value of *refinementWidth* depends on the degree of irregularity of the recording^3^.

### 1.2 Comb algorithm pseudocode

A pseudocode is given in **Algorithm 1**, using Matlab-like notation. Therefore, the expression *x* : *y* is the vector [*x, x* + 1, …, *y*], and *x* : *y* : *z* is 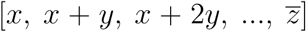, where 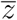 is the largest number lower or equal to *z*. Furthermore, the expression *a*(*b*) where *a, b* are vectors corresponds to those elements in *a* that are at positions specified in *b*.

#### Algorithm 1: Comb algorithm pseudocode.

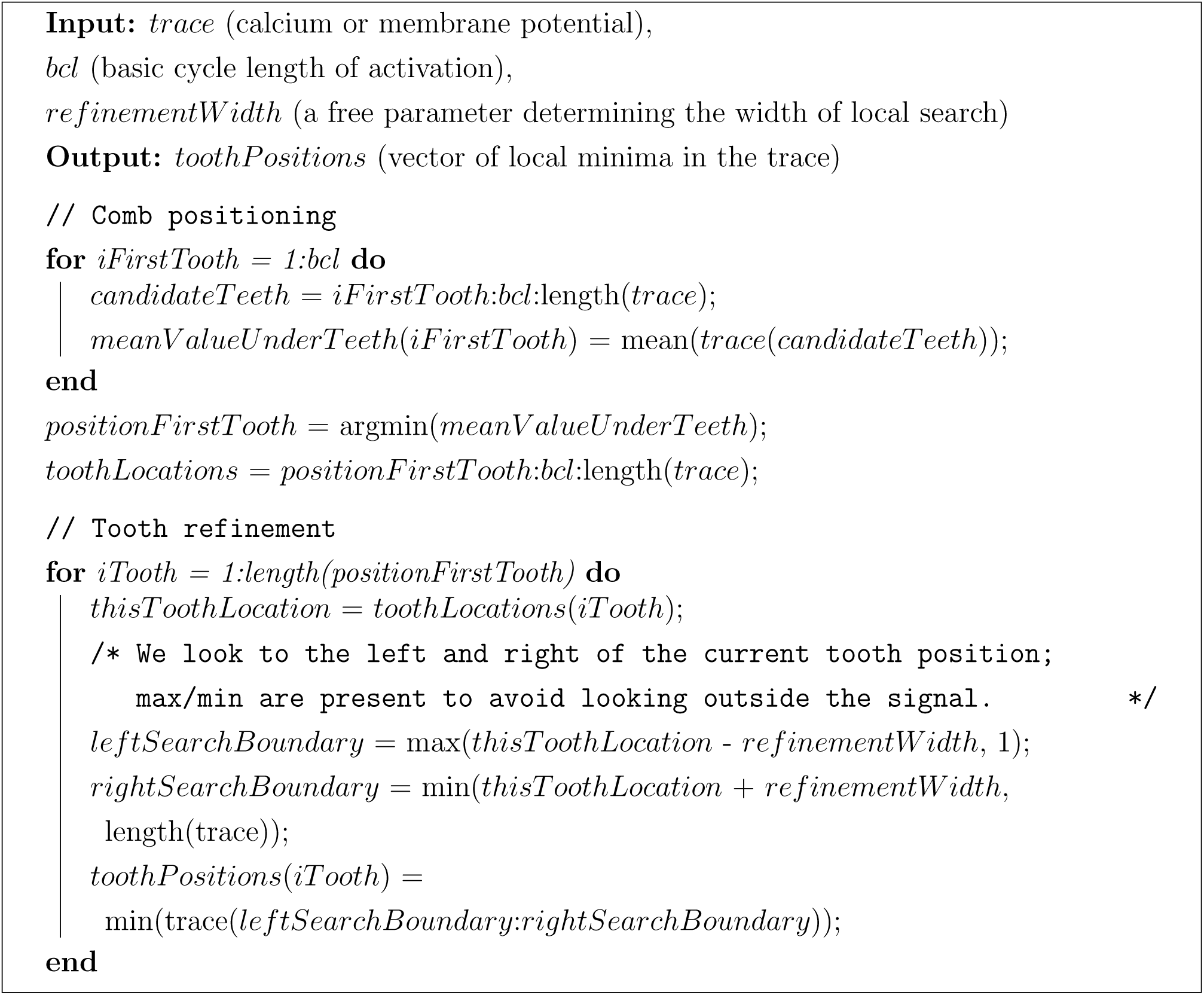

### 1.3 Algorithm adaptation for membrane potential recordings

Whereas in calcium imaging data, the local minima generally correspond to boundaries between calcium transients, this may not hold in optical or electrophysiological recordings of membrane potential, where an action potential is followed by diastolic interval where the signal is relatively flat (**Figure 2**). A local minimum may lie relatively anywhere within the diastolic interval, which complicates the optimal comb placement. For such case, we suggest the following:

**Figure 2:**
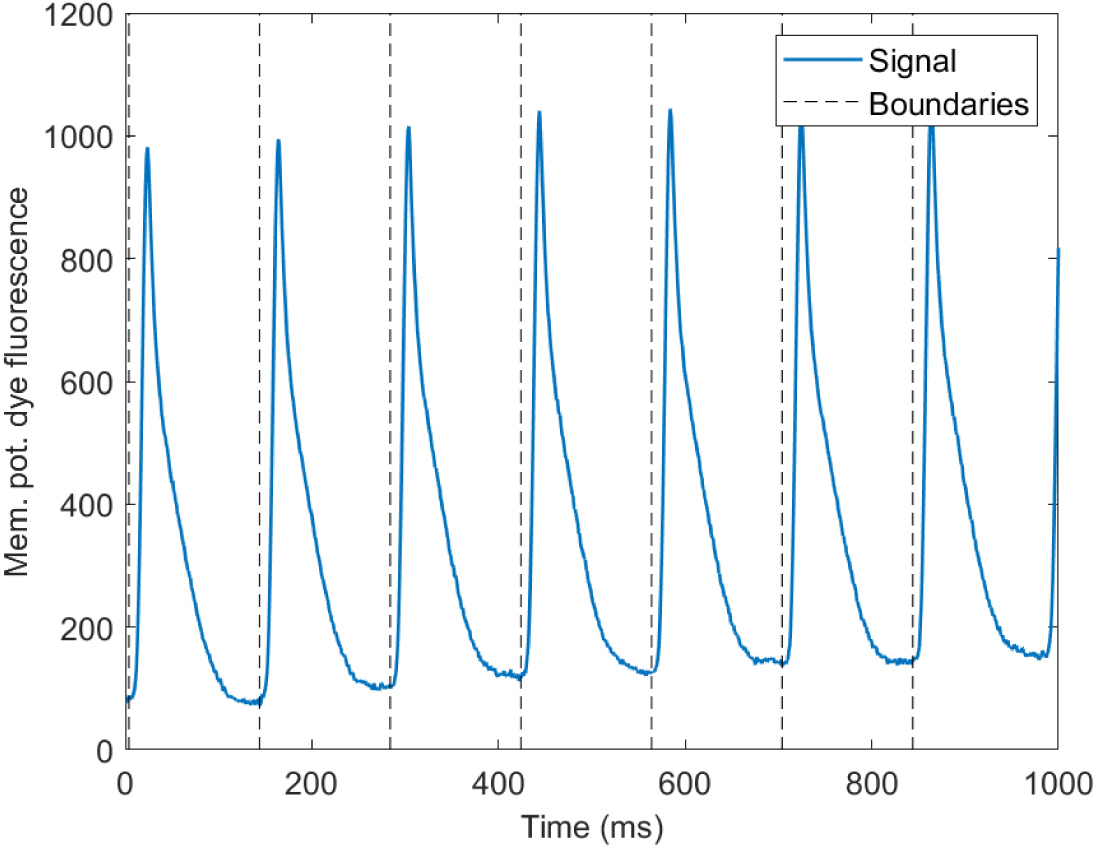
Membrane potential recording illustration. Sample recording of rat action potentials with boundaries between these highlighted. In larger mammals, the flat diastolic intervals are typically longer than action potentials themselves.

1. Flip the signal so that the peaks of action potentials point down.
2. Find minima using the baseline algorithm (the minima now correspond to peaks of action potentials).
3. Assume a sensible maximum duration of action potential upstroke (e.g., 20-30 ms) and place boundaries this long before the flipped signal minima. Therefore, one obtains signal boundaries that start right before the start of an action potential (**Figure 2**).

One exception when this approach could fail is extremely rapid pacing, where the diastolic interval is virtually nonexistent. In such a case, putting an action potential boundary 30 ms before an action potential peak could, in theory, hit the repolarization phase of the previous action potential. However, when the diastolic interval is extremely short, the basic version of the algorithm may be used instead.

Another approach to detection of action potentials may be to use the comb on upside-down flipped vector of derivatives of the signal; in this case, the comb position corresponds to the locations of action potential upstrokes.

## 2 Algorithm demonstration

Here are shown several examples of the algorithm’s performance, demonstrating how various signal artefacts are tackled. The experimental data were collected in a previous study in whole-heart imaging of rat Langendorff hearts [1].

### 2.1 Linear and nonlinear baseline drift, steps, and bleaching

Signal drift is a common finding in experimental recordings. It can result from gradual dye loading (leading to an increase in signal baseline over time), dye bleaching (reducing the signal baseline and amplitude over time), or more prosaic events such as the researcher bumping into the table with the imaging setup, or a droplet of solution passing over the imaged preparation, causing a transient change in signal intensity. The nonlinear drift is especially problematic for simpler algorithms for detection of calcium transients, given that it renders thresholding nearly useless unless the drift can be automatically learned and subtracted.

Both linear and nonlinear drift are processed correctly by the comb algorithm (**Figure 3A,B**). In addition, we tested the performance on very abrupt step-like change in the signal baseline; this arises during optogenetic experiments, when the preparation is illuminated by stepwise pulses [2, 3]. The algorithm functions well even when the step boundary falls within a calcium transient (**Figure 3C**).

**Figure 3:**
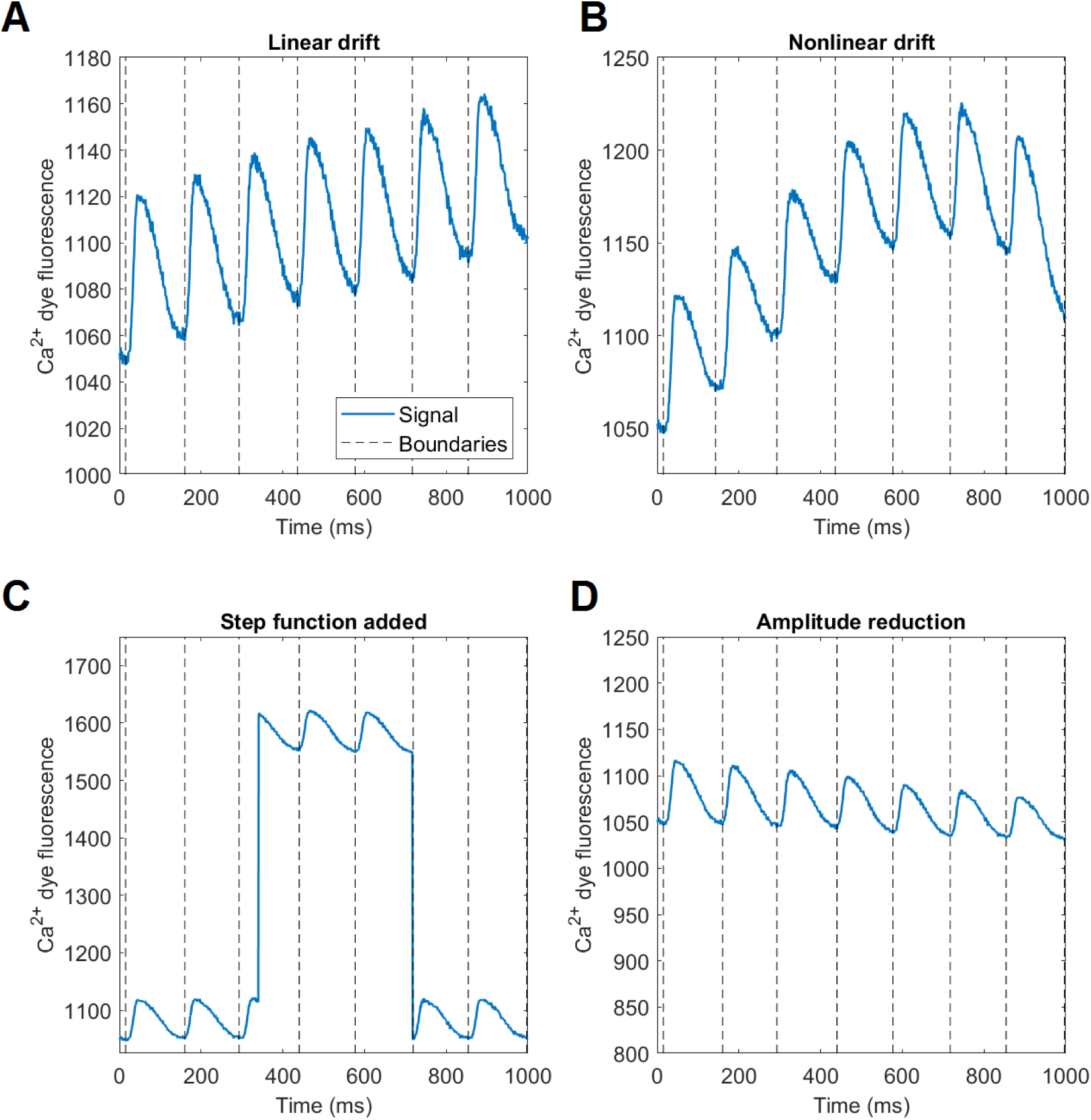
Drift artefacts. A) Linear baseline drift. B) Nonlinear baseline drift. C) Step-wise baseline perturbation. D) Baseline drift and amplitude reduction mimicking dye photobleaching.

Ultimately, we simulated photobleaching by reducing the baseline and amplitude, which did not present an obstacle to the algorithm (**Figure 3D**).

One type of drift-like behaviour which might prove problematic to the algorithm is periodic signal perturbation, e.g., arising from from contraction-induced motion artefacts in tissue mapping. In such a case, it is possible that there are signal minima that reflect tissue contraction rather than starts of cellular activation. Given that the frequency of contraction will be identical to the frequency of tissue activation, the artefact-induced minima may be detected as boundaries by the comb algorithm.

### 2.2 Gaussian noise

Gaussian noise is frequently found in optical and electrophysiological recordings. It may confuse thresholding algorithms by producing falsely positive objects of short duration. Approaches based on upstroke velocity may be also confused if the noise amplitude is high enough to mimick a segment of upstroke. The comb algorithm presented here can deal with low to very high amount of noise (**Figure 4A-C**), which follows from the global nature of the approach. If the recording is interpreted as the sum of true signal plus zero-mean noise, the noise component under the comb teeth is averaged over the teeth. Given that the noise is zero-mean, the longer the recording, the more teeth are present, making the average converge to zero, reducing the impact of noise on tooth positioning. However, this holds predominantly for initial positioning of the comb; the fine-tuning via local search around the initial tooth positions is vulnerable to noise to a greater degree. Therefore, not using excessively wide local search window is advisable for high-noise scenarios.

**Figure 4:**
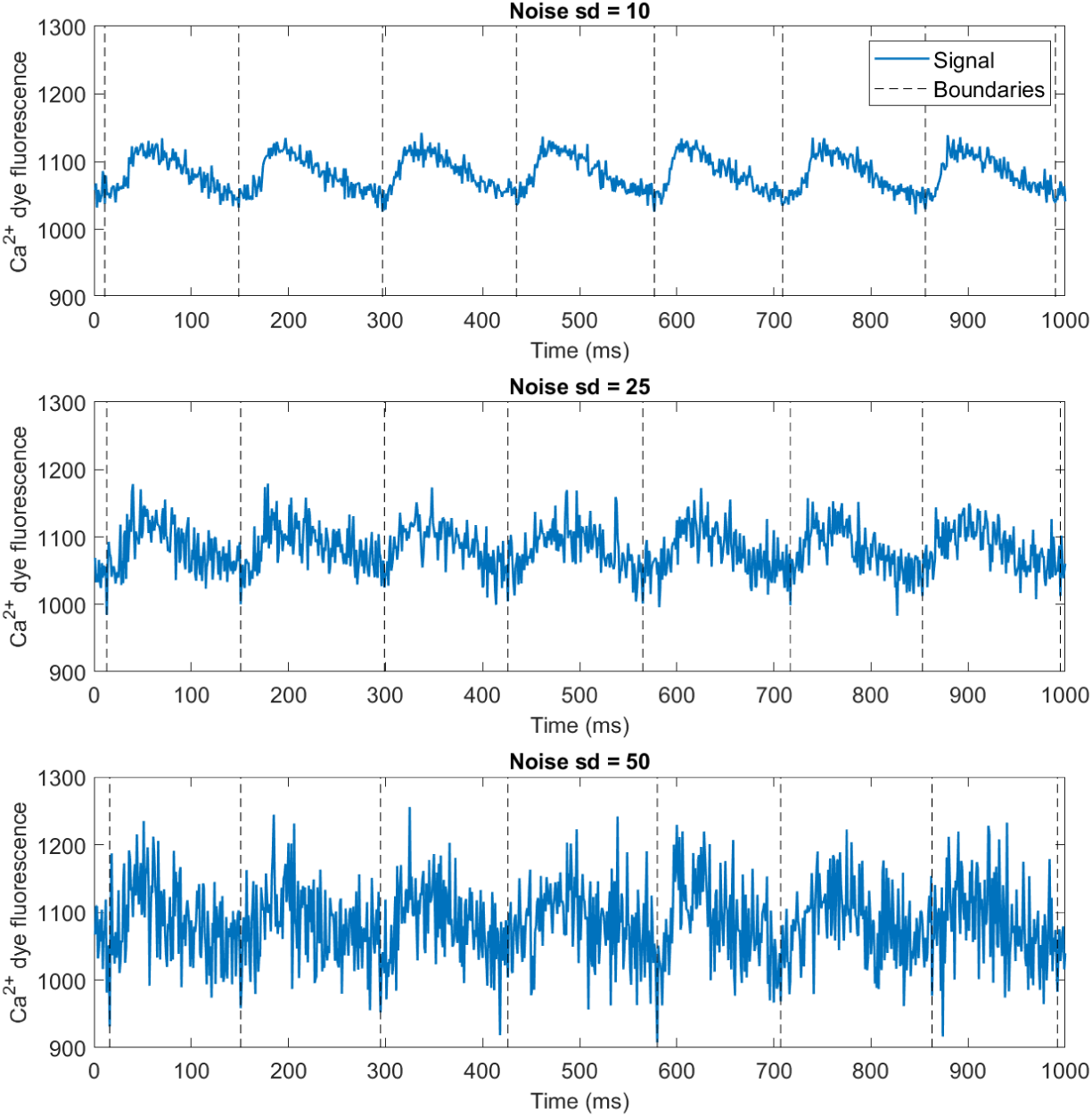
Gaussian noise. A-C) The calcium imaging recording from Figure 1 with Gaussian noise of standard deviation 10, 25, and 50 added, respectively.

### 2.3 Salt-and-pepper noise

Salt-and-pepper noise, also known as impulse noise, presents as sharp and sudden jump in a pixel’s intensity, which is typically sparse. This can be handled implicitly by the comb algorithm in many cases, such as **Figure 5A**. A case of salt-and-pepper which confuses the comb algorithm can be constructed, when a single extremely low value is present (**Figure 5B**). This acts as an attractor which forces the comb to put one of its teeth at this location. If such artefacts are present in data, the comb algorithm could be modified to minimize median of values under comb teeth, rather than the mean. Alternatively, median filtering may be applied to the signal, as it is usually an effective countermeasure against salt-and-pepper noise.

**Figure 5:**
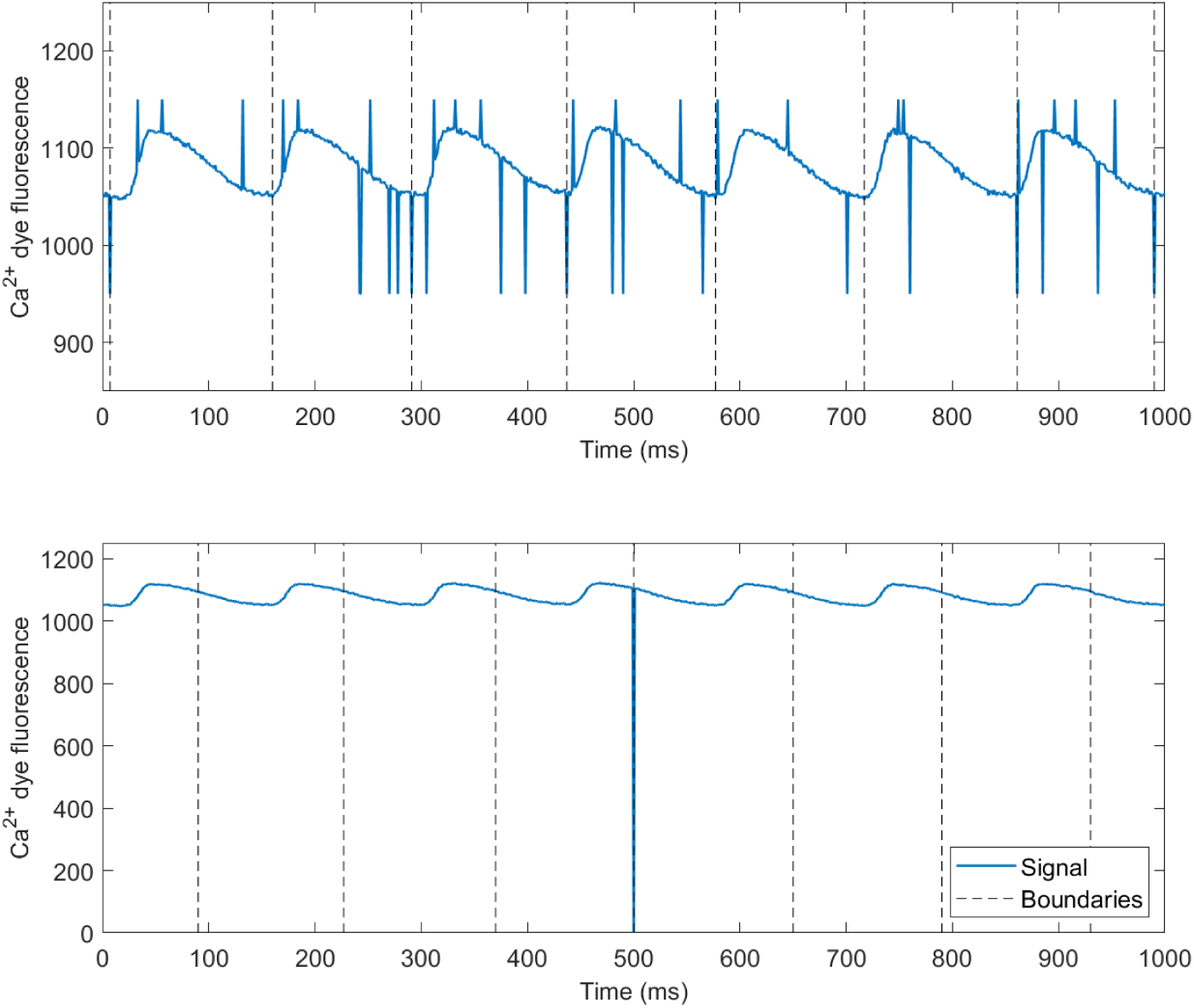
Salt-and-pepper noise. A) Sparse medium-amplitude salt-and-pepper noise. B) Extremely sparse large-amplitude salt-and-pepper noise.

### 2.4 Calcium alternans

One phenomenon which is not an artefact, but which may nevertheless prove troublesome, is calcium alternans, the periodic oscillation of large and small calcium transient at rapid pacing [4, 1]. The comb algorithm can easily detect separate transients when alternans is present (**Figure 6A**), even when it is severe (**Figure 6B**). Particularly the latter case of very severe alternans highlights why such data are hard to process e.g., via thresholding approaches, given that the range of thresholds which would detect the smaller calcium transients is narrow (and it may vary between different cells).

**Figure 6:**
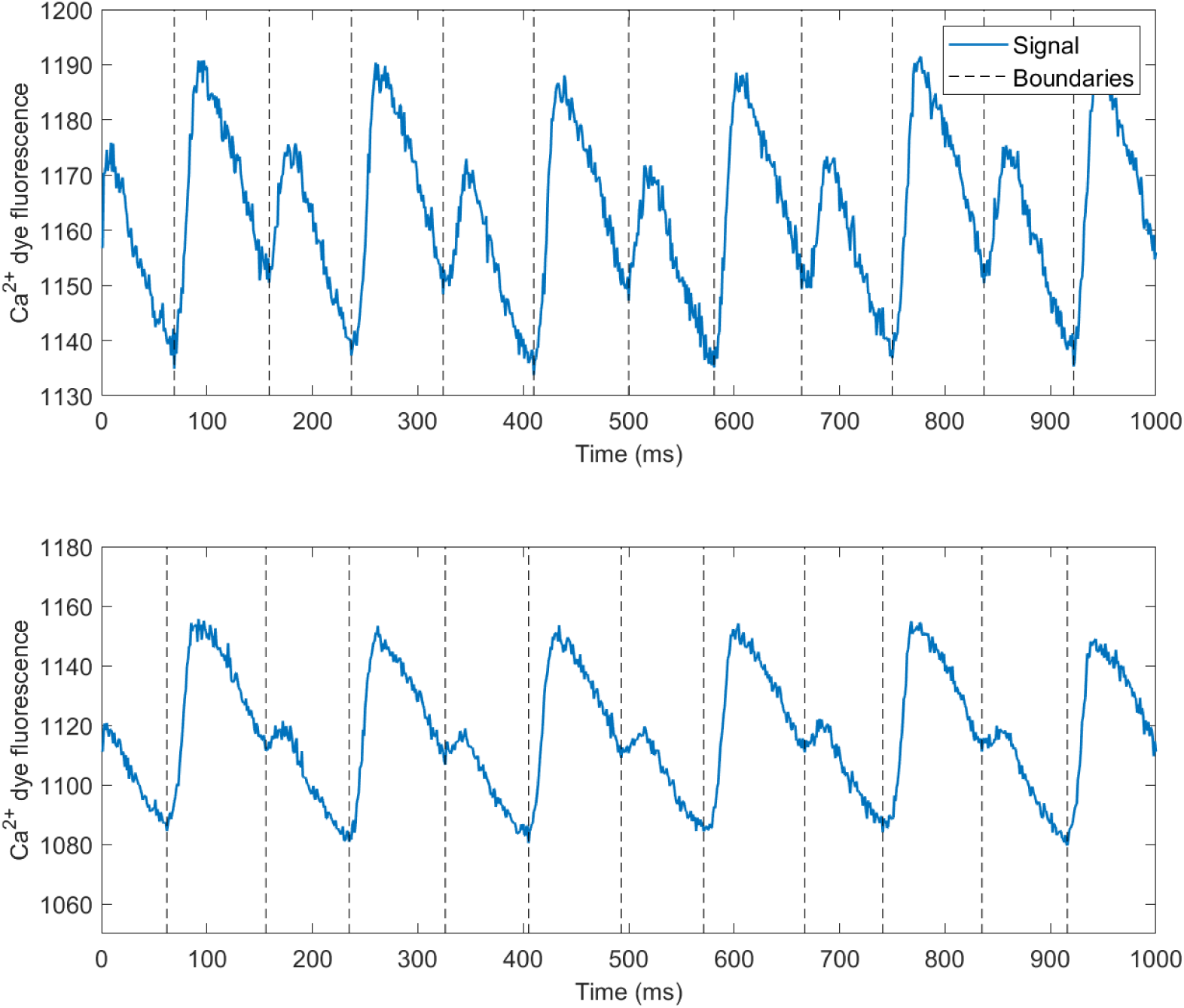
Calcium alternans. A) Medium-degree calcium alternans. B) very severe calcium alternans. Both cases were recorded at 85 ms bcl.

Particularly the case of severe alternans (**Figure 6B**) is one of motivations for the inclusion of the parameter allowing local search around initial comb tooth position. Given that the large calcium transients are longer than the short ones, a completely rigid comb with teeth 85 ms apart could not find transient boundaries accurately.

## 3 Discussion

Here we have presented a simple ‘comb’ algorithm for detection of calcium transients or action potentials in recordings of regularly activated cardiac preparations. The algorithm uses prior knowledge of the activation frequency to find all objects of interest at the same time, utilizing global information to overcome potential pitfalls in experimental data. It has the following advantages:

- It is resistant to noise and/or drift of signal, unlike simple alternative approaches based on thresholding or detection of rapid upstroke velocity.
- It does not need any reference data (like template matching), nor annotations (like machine learning). The independence on precise shape of the segmented object makes it readily applicable for other objects of interest (such as traces of ionic current measurements), as long as these have reasonably obvious minima or maxima.
- It is generally very simple, easy to implement in any language, also offering fast computational performance. In addition, it has only a single free parameter which furthermore has a very clear graphical intuition and is thus simple to set.

The fact that the algorithm requires an approximately fixed heart rate is a limitation, but it still covers a range of widely used scenarios in cardiac research. Fixed-rate pacing is often employed in experiments, e.g., to control for unwanted drug-induced changes in heart rate, or when alternans is elicited via the ‘S1’ protocol. In addition, the ‘S1-S2’ protocol used to study action potential duration restitution [5] can also be processed automatically. Furthermore, it is often the case that even an unpaced cardiac preparation has a stable-enough heart rate, making it suitable for processing by the comb algorithm. The accurate detection of action potential or calcium transient boundaries then greatly facilitates reliable extraction of biomarkers such as action potential duration, calcium transient duration, calcium transient amplitude (which then may be used to estimate the degree of calcium alternans), or area under a curve (e.g., calcium transient or ionic current measurement). The algorithm may be further adapted to achieve even greater flexibility:

- The basic algorithm uses equidistant comb teeth. However, the general approach can be used for any pattern of pacing as long as this is known in advance. E.g., a pacing protocol that employs pacing at gradually increasing rate corresponds to a comb with gradually reducing distance between teeth. Alternatively, alternans measurements typically employ pacing using blocks of fixed-rate stimuli, with the rate increasing between the blocks. This again may be fitted with a comb with corresponding blocks of teeth, where each block has equidistant teeth, but the distance is changed between blocks.
- The generation of non-equidistant comb teeth as in the previous point could be in theory automated via approaches detecting signal frequency over time. Additionally, ECG measurements are often employed as an additional feature in whole-heart recordings, which can be used to extract heart rate over time, and this could be used to generate an appropriate comb for a parallel imaging recording.

## Acknowledgment

I am very grateful to Prof. Gil Bub of McGill university for his suggestions to improving the manuscript.

## Footnotes

1 For clarity, we assume the recording is taken at 1 KHz equidistant sampling. The algorithm can be nevertheless naturally extended for any sampling rate.

2 It is not necessary to go beyond bcl ms, given that the resulting comb would be identical to a previously considered one.

3 In our experience, 5 ms is sufficient for generally regular recordings, while 15 ms is enough even for recordings with marked beat-to-beat variability.

